# A short cut to sample coverage standardization in meta-barcoding data provides new insights into land use effects on insect diversity

**DOI:** 10.1101/2025.01.27.635018

**Authors:** Mareike Kortmann, Anne Chao, Chun-Huo Chiu, Christoph Heibl, Oliver Mitesser, Jérôme Morinière, Vedran Bozicevic, Torsten Hothorn, Julia Rothacher, Jana Englmeier, Jörg Ewald, Ute Fricke, Cristina Ganuza, Maria Haensel, Christoph Moning, Sarah Redlich, Sandra Rojas-Botero, Cynthia Tobisch, Johannes Uhler, Jie Zhang, Ingolf Steffan-Dewenter, Jörg Müller

## Abstract

The use of metabarcoding for insect species identification has grown rapidly, but the absence of abundance data hinders meaningful diversity metrics like sample coverage-standardized species richness. Additionally, the vast number of taxa often lacks a unified phylogeny or trait database. We present a framework for constructing a phylogenetic tree encompassing the majority of insect families, standardisation of sample coverage (an objective measure of sample completeness) and assessment of both taxonomic and phylogenetic diversity using the Hill series for metabarcoding data. Applied to central Europe, our framework analysed insect diversity from 400 families along a land-use gradient. Results revealed land-use intensity significantly affects sample coverage, emphasizing the need for biodiversity standardization. After standardization, taxonomic diversity declined by 27–44%, and phylogenetic diversity by 13–29% across 39,000 Operational Taxonomic Units, from forests to agricultural areas. Rare species exhibited greater phylogenetic diversity loss than taxonomic, while dominant species showed smaller phylogenetic losses but stronger declines in taxonomic diversity. Our findings underscore agriculture’s detrimental effects on specific insect taxa, even after adjusting for sample coverage, and provide new insights into the loss of functional diversity, as represented by phylogeny.

## Introduction

Evidence of declining insect populations has alarmed both the scientific community and society. The complex interactions among various factors potentially responsible for the decline in different biodiversity metrics of insect communities continue to fuel controversial debates (Cardoso et al., 2020; Desquilbet et al., 2021; Saunders et al., 2020).

During the last decade, several studies have analysed the temporal trends of insect declines in different regions of the world. One of the most prominent studies by Hallmann et al. (2017) showed a drastic decline in insect biomass in protected areas. Neff et al. (2022) also observed that climate and land-use changes affect the decline of insect species distributions in butterflies, grasshoppers, and dragonflies. Similarly, a temporal analysis of bumblebees in Europe showed that suitable habitat will likely be reduced by at least 30% due to climate change and land use (Ghisbain et al., 2024). However, trends in insect populations are highly heterogeneous among different taxa and environments and depend on the target metric like biomass, abundance or diversity measures (Macgregor et al., 2021; Pilotto et al., 2020; van Klink et al., 2024). Recent studies showed the importance of considering the different characteristics of insect communities to gain a better understanding of the ongoing changes in insect populations. For example, van Klink et al. (2024) showed that the most severe temporal decline in the abundance of terrestrial insect communities occurs in dominant and not rare species.

## Introducing a framework for comprehensive diversity measures

One significant challenge in studying insect decline stems from their vast diversity. To efficiently cover such a broad range of taxa, metabarcoding has become increasingly prevalent. However, due to the lack of reliable abundance data equivalent to individual counts from sequencing, most studies rely solely on observed species numbers (often incorrectly named as species richness (see Gotelli & Colwell 2001), neglecting frequency distributions. This approach imposes two distinct limitations: the inability to standardize for varying levels of sample coverage and the inability to analyse more meaningful diversity metrics, such as species richness (Gotelli & Colwell, 2001), beyond mere species counts. Moreover, the sequence data obtained through metabarcoding often remains underutilized, despite its potential to provide valuable insights into phylogenetic relationships.

Sampling of insects or other diverse groups usually leaves some species undetected. Hence, insect samples are generally incomplete (Chao et al., 2014). But the degree of completeness can vary greatly and systematically between habitat types, sampling methods and species communities. Therefore, scientists have developed frameworks to calculate and control for sample coverage (or simply coverage), an objective measure of sample completeness. Coverage is defined as the proportion of individuals or frequencies in the entire community belonging to species detected in a sample.

We have set up a framework that allows us to consider taxonomic and phylogenetic diversity, but also the variation in sample coverage across habitats, an aspect that has been neglected by most previous analyses of insect diversity trends (but see van Klink (2024)). Contrary to most people’s intuition, Alan Turing showed that sample coverage can be very accurately estimated based on the number of singletons in a sample (Chao & Jost, 2012). This usually requires data on species frequency distributions, which is still an open area of research when dealing with metabarcoding data (Chiu & Chao, 2016; Elbrecht et al., 2017; Pont et al., 2023). However, we can use the number of reads generated during sequencing to describe the frequency distribution of species or other taxonomic units within a sample. This is possible after estimating the true number of singletons in the sample based on Turing’s wisdom (Chao et al., 2020; Chao & Jost, 2012) on statistical methods (Chiu & Chao, 2016). Within our framework, an adapted method based on Chiu and Chao (2016) filters highly diverse metabarcoding data to use the estimated true number of singletons in a sample. This now allows us to use the frequency of reads to describe species distributions within samples and to calculate sample coverage for metabarcoding samples (Chiu & Chao, 2016).

Nevertheless, even after controlling for sample coverage, we still have different species frequency distributions within our observed communities or samples. To incorporate species frequency distributions into biodiversity measures, a consensus has emerged among ecologists in biodiversity research that Hill numbers (Hill, 1973) (effective number of species) should be used to quantify species or taxonomic diversity ((Ellison, 2010) and subsequent papers). Hill numbers allow us to calculate diversity values with focus on rare (q=0 or species richness), common (q=1 or Shannon diversity) and dominant species (q=2 or Simpson diversity).

It is important to note that the Hill numbers in this study are based on metabarcoding reads, which reflect a combination of the number of individuals sampled, their biomass, and taxon-specific amplification biases, even after controlling for differences in insect size through sieving (see Methods). Still, metabarcoding reads can be used as an approximation for abundances when analysing entire samples and not making taxon-specific comparisons. Additionally, since we calculated sample coverage within each sample, we used raw read counts rather than relative read counts. Chiu and Chao (2016) established the validity of using such measures for microbes, which face similar challenges in quantifying abundance. To avoid ambiguity, we will refer to abundance distributions as frequency distributions.

In addition to analysing taxonomic diversity, functional diversity serves as a widely accepted metric to better understand ecosystem functioning and resilience, (Bregman et al., 2015). However, due to the absence of a consistent method or database for analysing functional traits across all terrestrial insect families (Moretti et al., 2017), we employed phylogenies as a practical proxy (Cadotte et al., 2012). To date, no comprehensive phylogeny exists that encompasses the majority of insect families and resolves to the species or OTU level, limiting the analysis of metabarcoding data. To address this, we developed a workflow to calculate a phylogenetic tree for large numbers of Operational Taxonomic Units (OTUs) in metabarcoding samples. This workflow is based on a backbone tree for insect families combined with the genetic sequences provided by metabarcoding. The backbone tree, which reflects deep evolutionary lineages (Rainford et al., 2014), serves as a foundational framework, while CO1 sequences from metabarcoding provide finer resolution within insect families. Although the backbone tree by Rainford et al. (2014) is the most comprehensive for insect families to date, it can easily be replaced by newer or improved versions in our workflow. Additionally, our method is not restricted to CO1 sequences but is flexible enough to incorporate other sequence types, such as ITS sequences in fungi. This combined approach introduces a novel method for calculating phylogenetic and taxonomic diversity measures in hyperdiverse insect samples. Importantly, it standardizes comparisons across samples with varying coverage, facilitating the analysis of species richness (q0), Shannon diversity (q1), and Simpson diversity (q2). Within this unifying framework of Hill numbers, we can also directly compare diversity measures for all numbers of q and between taxonomic and phylogenetic diversity.

## Analysing impacts of land use, climate and weather on insect communities

We used this framework to quantify the effect of major land use types on overall insect diversity in Central Europe. For the first time we can consider the different requirements for a mathematically correct estimation of diversity from large bulk samples based on metabarcoding (Chao & Jost, 2012; Gotelli & Colwell, 2001). In specific, we disentangle the effect of different land use intensities at regional and local scale, controlled for climate (long-term weather over 30 years) and weather (day-to-day atmospheric conditions) using the LandKlif study design which is located in a cultural landscape in Bavaria (Germany). It covers five climatic zones, four different local land-use types (forest, grassland, arable land and settlement) and three regional land-use types (semi-natural, agricultural and urban) with increasing land-use intensity (Redlich et al., 2022). From these study plots, we analysed 1,259 Malaise trap insect samples, containing 41,489 Operational Taxonomic Units (OTUs) of insects, of which 39,113 OTUs were assigned to 400 different insect families. Malaise traps capture the broadest range of insect taxa in terms of species and families. Although they primarily focus on flying insects, they provide a valuable link between traps that target ground-dwelling insects, such as pitfall traps, and those designed for flying insects, such as flight interception traps. Since Malaise traps collect a wide spectrum of Diptera and Hymenoptera (Fig. 1), which are difficult to identify visually, they are often paired with metabarcoding for species identification.

**Fig. 1.**
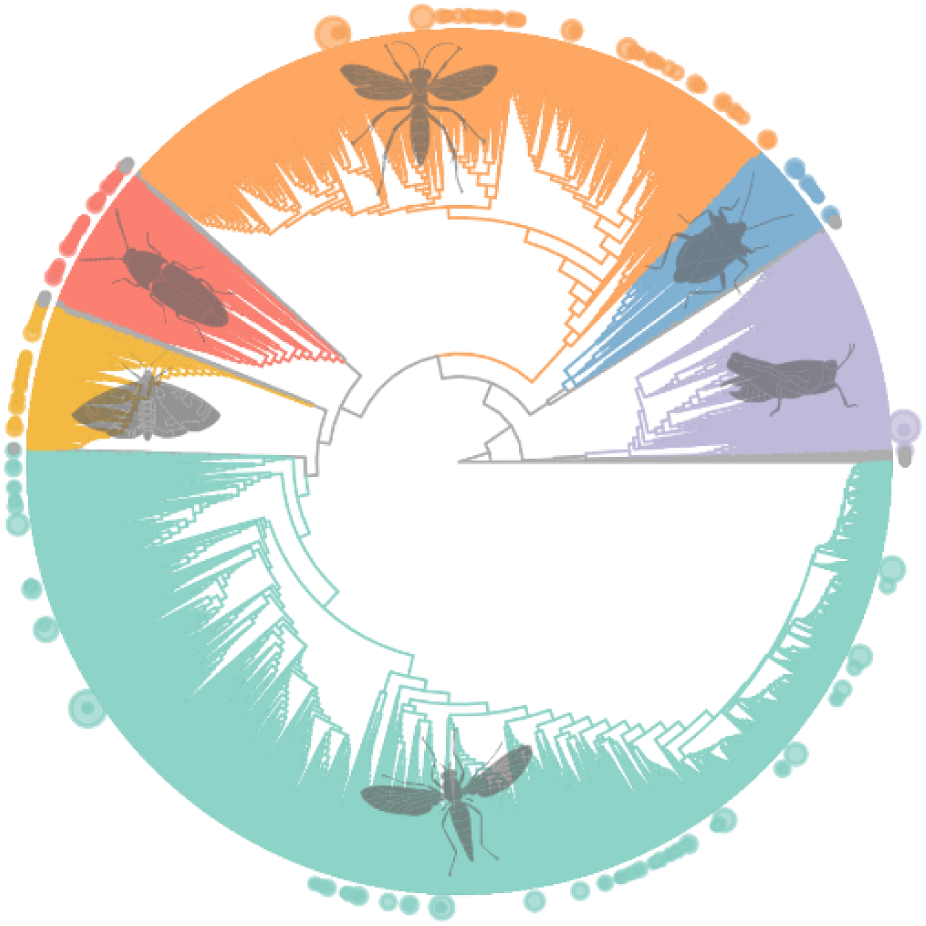
Phylogenetic tree of 39k OTUs from 400 insect families and their diversities. The main insect orders are depicted in colour (in a clockwise direction Hymenoptera in orange, Hemiptera in blue, Orthoptera in lilac, Diptera in aquamarine, Lepidoptera in yellow and Coleoptera in red). The remaining smaller orders are depicted in grey. The dots at the end of the branches represent families, with the dot size indicating the number of Operational Taxonomic Units within each family. The tree was calibrated with fossil calibration points from Rainford et al. (2014).

## Material and methods

### Study area

We based this study on data from the LandKlif project (https://www.landklif.biozentrum.uni-wuerzburg.de), in Bavaria, Germany. A detailed description of the study design can be found in Redlich et al. (2022). Based on a grid cell system covering all of Bavaria, we selected 60 grid cells (quadrants), creating independent gradients of land use and climate. We based regional land-use types at the quadrant level, classifying them into three categories based on Corine land cover data from 2012: Semi-natural regions were dominated by natural vegetation and forest; agricultural regions had at least 40% arable land; and urban regions comprised >14% urban areas. Therefore, each regional land-use type contained a higher proportion of each respective land use than the region-wide average. Given that quadrants were classified in three land-use types and five climate zones (using the multi-annual mean temperature of the period 1981-2010; DWD citation in Redlich et al. (2022)), 60 quadrants were selected to have 15 climate-land-use combinations, each with four replicates. Within each quadrant, three survey plots were selected based on the three most common land-use types of the quadrant. The local land-use types considered in this study were forest, grassland, arable land, and settlement. This resulted in 179 plots with a spatial extent of up to 480 km^2^.

### Insect sampling

Detailed information on insect sampling can be found in Uhler et al. (2021). We installed one Malaise trap at each plot centre. Malaise traps were based on the Townes Malaise trap model with a black roof and slightly smaller size (Uhler et al., 2022). We used ethanol (80%) as the trapping fluid to ensure preservation of DNA for barcoding. Traps were activated from mid-April to mid-August and emptied every two weeks, resulting in eight sampling campaigns and a total of 1259 insect samples. We separated the samples into smaller and larger insects using an 8 mm sieve to improve barcoding results. Separation into smaller and larger individuals also allows to control for differences in biomass, improving the role of reads as a surrogate for species frequency distribution within samples. Species were identified using CO1-5P (mitochondrial cytochrome oxidase 1) DNA metabarcoding following the laboratory and bioinformatics pipelines reported in Hausmann et al. (2020), which results in Operational Taxonomic Units (for most Orders the mean OTU number per plausible species ranges between 1 and 2, (Buchner et al., 2024)) and associated read counts.

Climate, local temperature, and humidity.

We recorded local temperature and humidity (collectively called ‘weather’) with ibutton thermologgers (type DS1923). At each site, we mounted one datalogger on a wooden pole, protected from direct sun exposure. We averaged hourly measurements of air temperature and relative humidity across the sampling periods. We calculated annual mean temperature and precipitation for each plot (from 1981-2010) based on gridded monthly datasets provided by the German Meteorological Service (DWD) with a horizontal resolution of 1 km (collectively called ‘climate’).

### Statistical analysis

We used 1293 insect samples for analyses of biodiversity and insect communities. We conducted all analyses in R 4.3.1 (R Core Team, 2023). We present an overview of the statistical methods in Supplement 1, Figure S1.1 with references to the R scripts (Zenodo repository https://doi.org/10.5281/zenodo.14747928).

### Singleton filter

We first created a sample-OTU-read-matrix and used a procedure based on a method from Chiu and Chao (2016) to filter unrealistic singletons and therefore control for sequencing errors (Fig. S1.1). This method uses a nonparametric estimator of the true singleton count of each sample. In turn, we based the estimate of the true singleton counts on the number of doubletons, tripletons and quadrupletons. From each sample, we removed the number of surplus singletons by random selection. We made slight improvements to the singleton filter originally proposed by Chiu and Chao (2016). The enhanced version can be found in R script 1. To control for differences between randomly filtered datasets, we repeated the random selection and our analyses of phylogenetic diversity and community matrices five times. Since the results of the different MRMs showed only very little variation, and estimates and p-values of the models of phylogenetic diversity showed no differences, we kept just one matrix for our results.

### Phylogenetic tree

We built a phylogenetic tree based on metabarcoding sequences and a backbone tree by Rainford et al. (2014). Metabarcoding sequences are well suited to differentiate species, but they are not able to explain older phylogenetic divisions like the separation of different orders. To gain reliable information on older lineages, we used a dated phylogenetic tree of insects that covers relations down to families as backbone tree (Rainford et al., 2014). We added missing families to the backbone tree, but only when we had sufficient information about the sister families. To resolve relations within families, we used CO1 sequences used for metabarcoding. Sequences were aligned within each family using the *AlignSeqs* function from the *DECIPHER* package (Wright, 2015). We calculated distance matrices for each alignment with *DistanceMatrix* function from *DECIPHER* and estimated phylogenetic trees with the *TreeLine* function using the Neighbour-Joining method. The estimation of subtrees can be found in R script 3. The family subtrees were then added to the backbone tree at the respective family node with the *bind.tree* function from the *ape* package (Paradis & Schliep, 2019). Due to the ultrametric structure of the backbone tree, family ages were then all the same. To calibrate all branch length, we used *make_bladj_tree* from datelife package, which uses the *bladj* function from *phylocom* (Webb et al., 2008). As calibration points we used the fossil data from Rainford (Rainford et al., 2014). To make the calibrated tree ultrametric, we used the *forceEqualTipHeights* function from the *ips* package. The assembly of the final tree can be found in R script 4 and the calibration in script 5.

### Multiple regression of distance matrices

We calculated community distances for sample coverage standardized communities (SC = 0.996) along the Hill numbers for rare, common and dominant species based on Jaccard, Horn and Morisita-Horn indices using an adapted function based on the inext.beta3D function (R script 09 and 10), an approach by Chao et al. (2023). We calculated Euclidean distances for the mean day of sampling, climate and weather variables and geographic location (coordinates) using the *dist* function in R. Gower distances were calculated for local and regional land-use types using the *daisy* function from the *cluster* package (Gower, 1971; Maechler et al., 2022). Calculations of the distance matrices of environmental parameters can be found in R script 11. We calculated multiple regressions on distance matrices with the *MRM* function from the *ecodist* package (Goslee & Urban, 2007) to test for effects of distances in land use and climate and weather on community distances (R script 12).

### Calculation of taxonomic and phylogenetic diversity

To calculate biodiversity measures, we used the *iNEXT.3D* package in R (Chao et al., 2021). We used the *iNEXT3D* function to calculate the observed taxonomic and phylogenetic diversity as well as the sample coverage of each insect sample (R script 6).

Coverage-based standardization is a mathematically elegant and statistically robust way to standardize samples, and has been increasingly used in the ecological literature (Roswell et al., 2021). Based on our data, sample coverage in 179 plots ranged from 0.981 to 0.999 (Fig. S1.2). For highly diverse data, a small percentage of coverage, such as 2%, may contain many undetected species. Classic sample completeness is defined as the observed richness divided by the estimated true species richness. Since true richness typically cannot be accurately estimated for species data, classical sample completeness cannot be accurately estimated from the sample itself. In contrast, sample coverage, originally developed by the founder of computer science, Alan Turing, can be accurately estimated from the sample itself. For example, if we assume that we have three dominant species in a community, their proportions of individuals in the total assemblage are 50%, 30% and 15%, respectively, and there are 100 other species, each representing only 0.05%. If we take three observations and assume that three dominant species are observed, the sample coverage of this sample would be 95%, whereas the classical sample completeness would be 3/103 ∼ 3%. A very small percentage of coverage can contain many species due to the possible presence of vanishingly rare species. In this example, 5% coverage includes 100 species, whereas 95% includes only three species. Therefore, whether a coverage value is "relatively" high depends on the distribution of species abundance or reads. In our case, 98% coverage is relatively low.

To calculate the estimated taxonomic and phylogenetic diversity we used the *estimate3D* function. The *estimate3D* function allows the standardisation of diversity measures to a given sample coverage, thus controlling for differences in sample coverage. We chose a level of 99.6% sample coverage, the mean sample coverage in our data, to avoid diversity values that are mainly based on extrapolation. To test the robustness of our results, we additionally calculated all diversity measures for a standardized sample coverage of 98%, the lower range within our data. The results for a sample coverage of 98% are in Table S1.2.

We calculated taxonomic and phylogenetic diversity along Hill numbers q=0, q=1 and q=2 to consider relative read distributions. The interpretations of taxonomic diversity (Hill number) of orders q= 0, 1 and 2 are the following: for q=0 all species are counted equally without considering their relative frequency, hence diversity measures for q=0 are more sensitive to rare species. For q=1, each species is weighted in proportion to its frequency, which lays the focus on common or common species. For q=2, abundant or frequent species are weighted disproportionately, hence diversity measures for q=2 represent the effective number of very abundant or dominant species. Similar interpretations are valid for phylogenetic diversity of orders q = 0, 1 and 2 by replacing "species" with "lineages".

### Generalized additive models

Analyses of the diversity measures are based on models from Uhler et al. (2021). We used the *gam* function from the *mgcv* package to fit generalized additive models to test for the effects of land use and climate on the taxonomic and phylogenetic diversity along the Hill numbers for each sample. Taxonomic diversity values were rounded to integers to be equivalent to numbers of taxonomic units, and modelled with a negative binomial error term (R script 7). Phylogenetic diversity values were kept as decimal numbers and modelled with a Gaussian error term (R script 8). Predictors for land use included local land-use types (forest, grassland, arable land and settlement) and regional land-use types (semi-natural, agricultural and urban). The mean day of a trap-specific sampling period was modelled by a smoothed non-linear spline of time to account for seasonality, and an offset for sampling length to control for variation in individual sampling periods. A correlated plot-specific intercept (geographical location of the plot) was used to account for the spatial arrangement within and between grids and for repeated measurements per plot. Mean annual temperature and precipitation served as predictors of macroclimate. The mean local temperature and mean humidity at each plot were calculated for the specific sampling window of each insect sample.

Furthermore, we fitted generalized additive models to test for effects of land use and climate on sample coverage. The model included the same variables as mentioned above but a binomial error term. Significant differences between land-use categories were assessed by multiple post-hoc comparisons using the *glht* function from the *multcomp* package (Hothorn et al., 2008). Calculations of generalized additive models and post-hoc comparisons can be found in R scripts 7 and 8 for taxonomic and phylogenetic diversity, respectively. All model results for a standardized sample coverage of 9.6% are in Table S1.1. Additional results for a standardized sample coverage of 98% are in Table S1.2.

## Results

The results of the multiple regressions (MRM) show that the day of the year had the strongest effect on community composition (Regression coefficients: 0.046 (q0), 0.037 (q1), 0.025 (q2)) followed by local land-use types and climate and weather. Regional land use and geographic location had the smallest impact (Fig. 2). In addition, climate and day of the year (i.e. mean date from the timeframe at which the insects were sampled) had stronger effects on communities with a focus on rare (q0) and common species (q1).

**Fig. 2.**
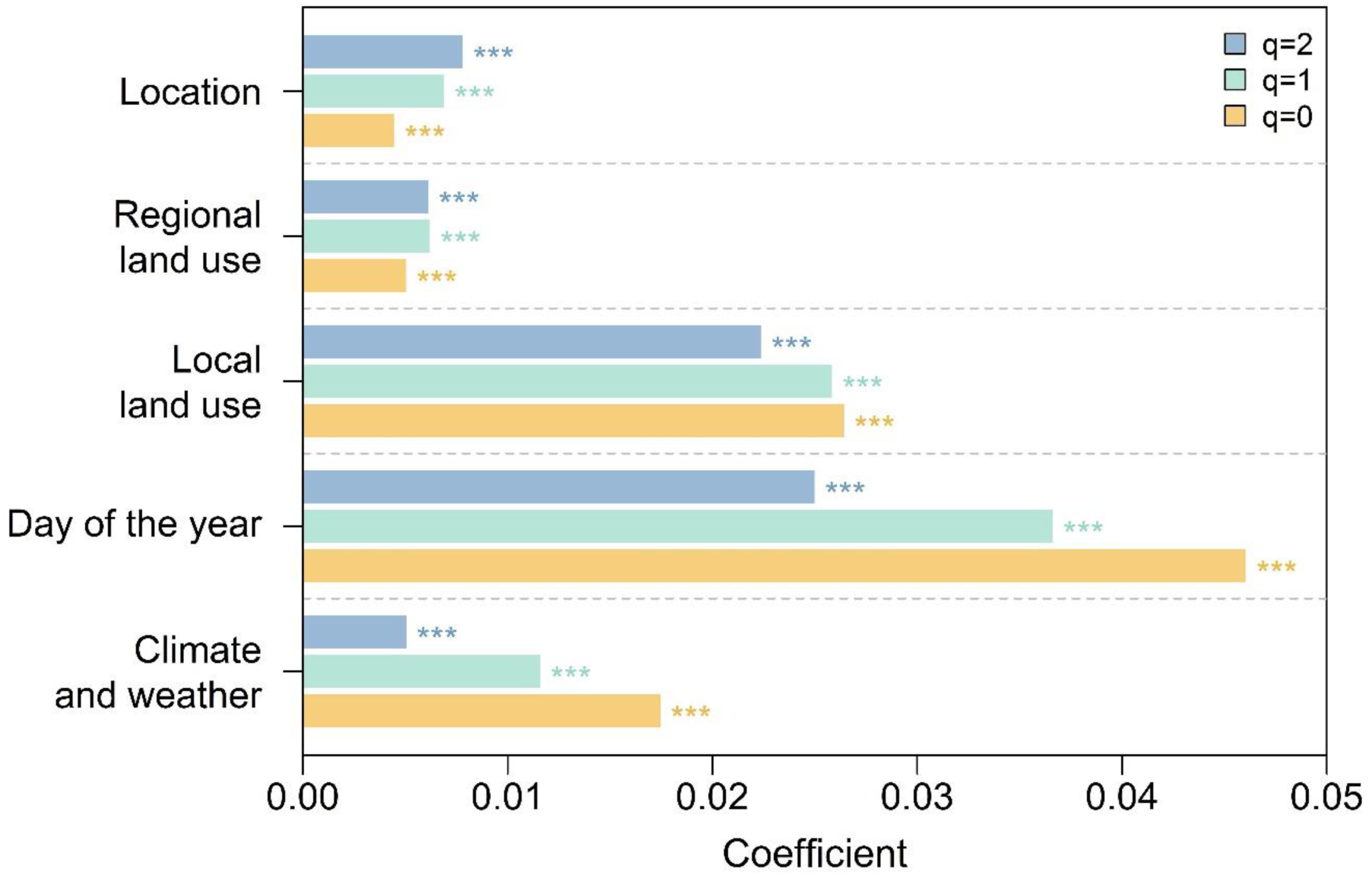
Model coefficients of multiple regressions on distance matrices, calculated with focus on rare (q=0), common (q=1) and dominant (q=2) species. Models included geographic distances (based on geographic coordinates in m), distances of regional and local land-use categories, day of the year and climate and weather. Models were calculated for coverage standardized dissimilarity matrices along the Hill numbers (q=0, q=1, q=2) to control for relative frequency distributions of reads generated during metabarcoding. Significant results (p ≤ 0.001) are marked with three stars.

Sample coverage was significantly higher in arable land compared to forests and in urban regions compared to semi-natural regions (Table S1.1). Sample coverage was also affected by the day of the year, with the lowest sample coverage occurring in July (Table S1.1 and Fig. S1.3).

Model results of the sample coverage-controlled diversity values showed that forests and semi-natural regions had the highest values of taxonomic (Fig. 3 a) and b)) and phylogenetic (Fig. 3 c) and d)) diversity. The negative effects of local and regional land use in comparison to forest and semi-natural environments were strongest for arable land and agricultural regions (for q1: Est.: -0.349, p < 0.001 and Est.: -0.154, p < 0.001).

**Fig. 3.**
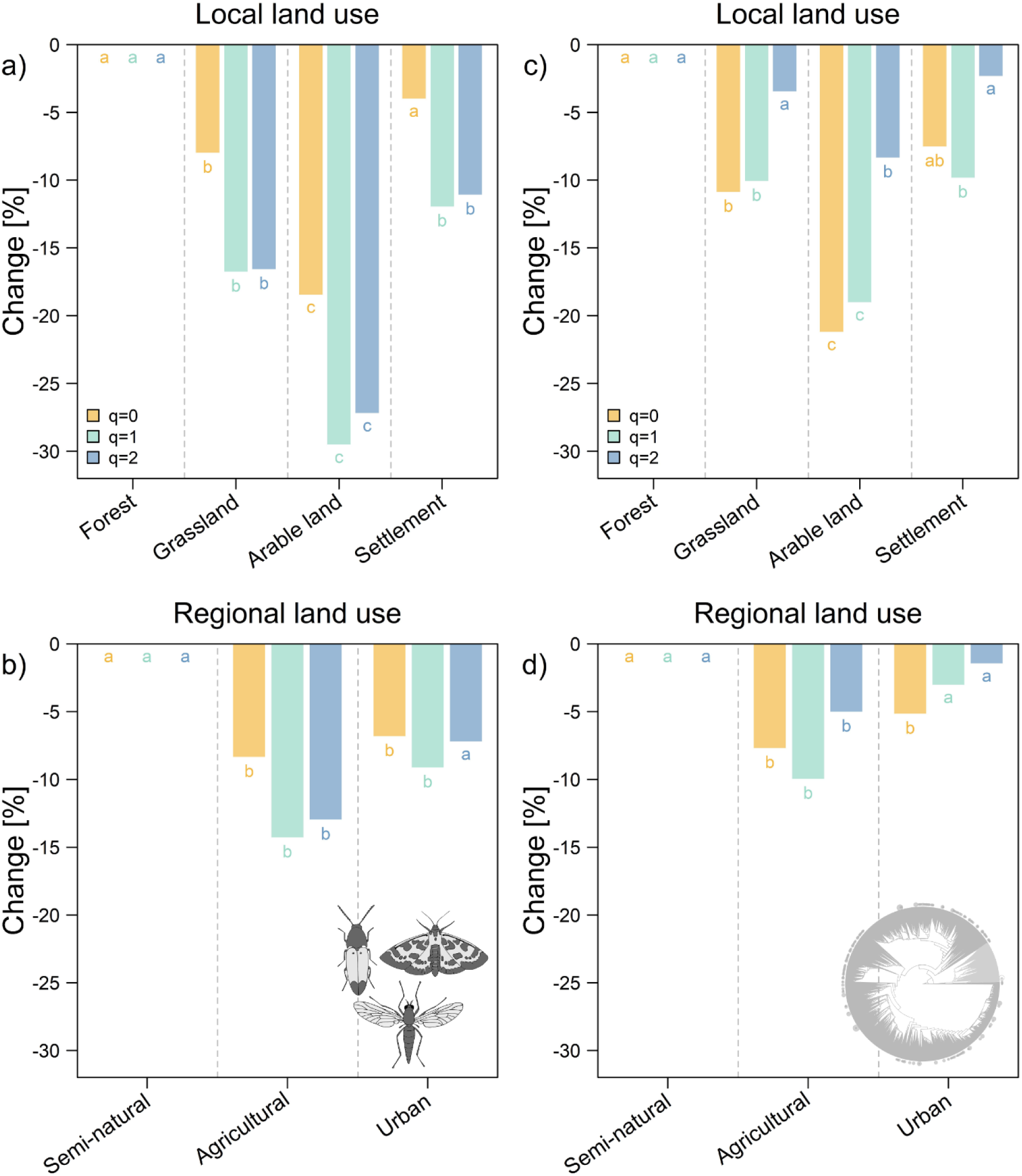
Multiplicative partial effects of land use on taxonomic and phylogenetic diversity based on generalized additive mixed models. Models included local and regional land-use categories and controlled for season, climate, weather and space. Taxonomic diversity values were calculated for a standardized sample coverage of 99.6%. The values are in comparison to (a) the local land-use type forest and (b) regional land-use type semi-natural. Models were calculated along the Hill numbers (q=0, q=1, q=2) to control for relative read distributions. We display all model parameters in Table S1. Letters indicate significant differences between categories (p < 0.05).

Within taxonomic diversity common (q1) and dominant (q2) insect species showed the strongest negative responses to land use. These effects occurred in grassland, arable land and settlements (Fig. 3 a)) as well as in agricultural regions (Fig. 3 b)). In contrast, taxonomic diversity in urban regions was only significantly reduced focusing on rare (q0) and common species (q1) (Fig. 3 b)).

The results for phylogenetic diversity followed the same patterns like taxonomic diversity regarding the different land-use types. Still, reductions in phylogenetic diversity were most pronounced for rare and common species (Fig. 3 c) and d)).

## Discussion

Using our new workflow, we demonstrated the impacts of land use on both taxonomic and phylogenetic insect diversity across the broad range of terrestrial insects captured by malaise traps, standardized for differences in sample coverage.

Sample coverage varied over the sampling period and was influenced by local and regional land-use type. The highest sampling coverage was observed in arable land and urban regions (Table S1.1), indicating these environments can be simplified. This suggests that with equivalent sampling efforts, a greater proportion of the insect communities can be effectively surveyed compared to more natural ecosystems. Conversely, the diversity observed in natural environments, such as forests, which generally exhibit lower sampling coverage, is underestimated relative to these simplified environments. Sample coverage was also affected by the day of the year (Table S1.1 and Fig. S1.3), with the lowest sample coverage occurring in July. During that time, we can also observe the highest biomass values, as shown in Uhler et al. (2021).

Furthermore, as shown in van Klink et al. (2024), sample coverage can also vary greatly between years, with an underestimation of diversity in years with lower numbers of individuals in the samples. Whereas it is no surprise that sample coverage can vary between seasons and years, these results show that we face two sources of sampling incompleteness in insect community studies: underestimation of diversity in species-rich habitats or landscapes, and underestimation within large samples, that contain huge amounts of biomass leading to increasingly undetected singletons (Chao & Jost, 2012). While further development of sequencing depth (Buchner et al., 2024) could increase the coverage within large samples, the bias in sample coverage caused by different environments will remain. Even conventional species-abundance data from selected Orders or Families (Seibold et al., 2019; van Klink et al., 2024) are faced by temporal variation in sample coverage (see findings in van Klink (van Klink et al., 2024)) and are affected by variation in sample coverage among habitats. We therefore strongly advocate standardisation of sample coverage, not only when comparing different habitat types, but also when comparing data within or between years, to avoid systematic sampling bias in diversity measures (Chao & Jost, 2012; Gotelli & Colwell, 2001).

Forests and semi-natural regions showed the highest values of taxonomic (Fig. 4 a) and b)) and phylogenetic (Fig. 4 c) and d)) diversity. The negative effects of local and regional land use were strongest for arable land and agricultural regions, with 18-30% and 8-14% less taxonomic insect diversity (OTUs), respectively, compared to forests. These results are in line with Uhler et al. (Uhler et al., 2021), who showed similar trends for Barcode Index Number (BIN) richness. It also confirms the general impression that agricultural areas have an impoverished biodiversity (Outhwaite et al., 2022; Raven & Wagner, 2021; Wagner, 2020), not only due to monotonous vegetation, but also due to negative effects of pesticides on non-target species (Nicholson et al., 2023).

## Common and dominant species react strongest in agricultural environments but not in urban regions

By considering different frequency distributions of species via the Hill numbers, we were able to show that common and dominant insect species showed the strongest negative responses to land use. These effects occurred in grassland, arable land and settlements as well as in agricultural regions (Fig. 3). Similar responses of highly abundant species were observed for population trends of hoverflies (Hallmann et al., 2021) and also for a broader selection of terrestrial insects over time (van Klink et al., 2024). These newly observed trends are in contrast to previous expectations of rare species or species with small range size being especially prone to population declines (Powney et al., 2019; Staab et al., 2023).

In contrast, taxonomic diversity in urban regions was only significantly reduced focusing on rare and common species (Fig. 3). Although urban environments generally have a negative impact on insect populations (Svenningsen et al., 2022; van Klink et al., 2020), e.g. through habitat loss due to sealing, they are still more heterogeneous and complex than agricultural areas (Fenoglio et al., 2021) and may therefore elicit more distinct responses from different taxa (Herrmann et al., 2023; Knop, 2016; Savage et al., 2015).

## Responses of phylogenetic diversity are driven by rare species

The results for phylogenetic diversity were similar to those for taxonomic diversity but more pronounced for rare and common species. We observed the greatest reduction in phylogenetic diversity in arable land for q0 and q1 (∼20%) (Fig. 4 c)). These findings suggest that species loss in agricultural areas does not occur randomly across the phylogeny. The weaker response of dominant species in terms of phylogenetic diversity indicates that larger lineages, such as Orders or Families, persist in these areas. In contrast, when focusing on species richness (q0) (i.e., all OTUs regardless of their abundance or frequency), we observe stronger effects on phylogenetic diversity, implying a non-random reduction in younger sections of the phylogenetic tree, such as genera. This aligns with a study on red-listed beetles, which also found a strong phylogenetic signal for the extinction risk of species (Seibold et al., 2015).

Focusing on common species (q1, Shannon diversity) we observed strong reductions in taxonomic and phylogenetic diversity. These results are quite alarming. Since a reduced phylogenetic diversity can also lead to less stability in a community (Cadotte et al., 2012), insect declines in agricultural environments might also accelerate in the future. The differences in Hill numbers also highlight the importance of considering read or frequency distributions in biodiversity analyses, since they can affect community responses and their extent of responses quite heavily.

## Conclusion

Our findings, which indicate that negative land-use effects on taxonomic diversity are more prominent in common and dominant species than in rare species, complement recent studies showing that temporal declines in insects are also most pronounced in dominant species (van Klink et al., 2024). However, while the aforementioned meta-analysis (van Klink et al., 2024) found that the temporal negative trends in diversity disappeared after standardization for sample coverage, the negative effects of more intensive land use in our study persisted after standardization.

In contrast, the negative effects on phylogenetic diversity were more pronounced for both rare and common species, underscoring the importance of incorporating phylogeny into conservation strategies. While this approach is widely accepted in the scientific community (Faith, 1992; Winter et al., 2013), the comprehensive inclusion of phylogenetic information in metabarcoding has been limited by the lack of available phylogenetic trees. Our new approach enables a straight forward integration of phylogenetic information in insect diversity studies with large metabarcoding datasets.

Taken together, our findings suggest that in temperate regions such as Central Europe, land use is the primary driver of overall insect diversity, whereas weather conditions mainly influence temporal population fluctuations, as indicated by changes in abundance and biomass (Müller et al., 2024; van Klink et al., 2024). However, a deeper understanding of how these two processes interact, including their impact on the extinction risk of individual species, is still lacking. To date, there are no long-term studies with climate and land-use gradients like those in this study, nor replicated land-use change experiments with appropriate controls (Outhwaite et al., 2022).

## Acknowledgements

We would like to thank Stefan Schmidt for his taxonomic support in this project and dedicate this article to his memory. Sadly, Stefan passed away during the course of this project. We would also like to thank Caryl S. Benjamin, Rebekka Riebl and Lars Uphus for their assistance in the field. We acknowledge funding from the Bavarian Ministry of Sciences and the Arts through the Bavarian Climate Research Network (bayklif) and the LandKlif project. MK was funded by the Bavarian Research Institute for Digital Transformation (bidt), an institute of the Bavarian Academy of Sciences, as part of the ROOT project (KON-22-024). JR and JMü acknowledge funding from the Bavarian State Ministry of Food, Agriculture and Forestry.

## References

1. Bregman, T. P., Lees, A. C., Seddon, N., Macgregor, H. E. A., Darski, B., Aleixo, A., Bonsall, M. B., & Tobias, J. A. (2015). Species interactions regulate the collapse of biodiversity and ecosystem function in tropical forest fragments. Ecology, 96(10), 2692–2704. 10.1890/14-1731.1

2. Buchner, D., Sinclair, J. S., Ayasse, M., Beermann, A., Buse, J., Dziock, F., Enss, J., Frenzel, M., Hörren, T., Yuanheng, L., Monaghan, M. T., Morkel, C., Müller, J., Pauls, S. U., Richter, R., Scharnweber, T., Sorg, M., Stoll, S., Twietmeyer, S., … Leese, F. (2024). Upscaling biodiversity monitoring : Metabarcoding estimates 31,846 insect species from Malaise traps across Germany. Molecular Ecology Resources.

3. Cadotte, M. W., Dinnage, R., & Tilman, D. (2012). Phylogenetic diversity promotes ecosystem stability. Ecology, *93*(8 SPEC. ISSUE), 223–233. 10.1890/11-0426.1

4. Cardoso, P., Barton, P. S., Birkhofer, K., Chichorro, F., Deacon, C., Fartmann, T., Fukushima, C. S., Gaigher, R., Habel, J. C., Hallmann, C. A., Hill, M. J., Hochkirch, A., Kwak, M. L., Mammola, S., Ari Noriega, J., Orfinger, A. B., Pedraza, F., Pryke, J. S., Roque, F. O., … Samways, M. J. (2020). Scientists’ warning to humanity on insect extinctions. Biological Conservation, 242(November 2019). 10.1016/j.biocon.2020.108426

5. Chao, A., Gotelli, N. J., Hsieh, T. C., Sander, E. L., Ma, K. H., Colwell, R. K., & Ellison, A. M. (2014). Rarefaction and extrapolation with Hill numbers: A framework for sampling and estimation in species diversity studies. Ecological Monographs, 84(1), 45–67. 10.1890/13-0133.1

6. Chao, A., Henderson, P. A., Chiu, C. H., Moyes, F., Hu, K. H., Dornelas, M., & Magurran, A. E. (2021). Measuring temporal change in alpha diversity: A framework integrating taxonomic, phylogenetic and functional diversity and the iNEXT.3D standardization. Methods in Ecology and Evolution, 12(10), 1926–1940. 10.1111/2041-210X.13682

7. Chao, A., & Jost, L. (2012). Coverage-based rarefaction and extrapolation: Standardizing samples by completeness rather than size. Ecology, 93(12), 2533–2547. 10.1890/11-1952.1

8. Chao, A., Kubota, Y., Zelený, D., Chiu, C. H., Li, C. F., Kusumoto, B., Yasuhara, M., Thorn, S., Wei, C. L., Costello, M. J., & Colwell, R. K. (2020). Quantifying sample completeness and comparing diversities among assemblages. Ecological Research, 35(2), 292–314. 10.1111/1440-1703.12102

9. Chao, A., Thorn, S., Chiu, C. H., Moyes, F., Hu, K. H., Chazdon, R. L., Wu, J., Magnago, L. F. S., Dornelas, M., Zelený, D., Colwell, R. K., & Magurran, A. E. (2023). Rarefaction and extrapolation with beta diversity under a framework of Hill numbers: The iNEXT.beta3D standardization. Ecological Monographs, 93, e1588. 10.1002/ecm.1588

10. Chiu, C. H., & Chao, A. (2016). Estimating and comparing microbial diversity in the presence of sequencing errors. PeerJ, 4, e1634. 10.7717/peerj.1634

11. Desquilbet, M., Cornillon, P. A., Gaume, L., & Bonmatin, J. M. (2021). Adequate statistical modelling and data selection are essential when analysing abundance and diversity trends. Nature Ecology and Evolution, 5(5), 592–594. 10.1038/s41559-021-01427-x

12. Elbrecht, V., Peinert, B., & Leese, F. (2017). Sorting things out: Assessing effects of unequal specimen biomass on DNA metabarcoding. Ecology and Evolution, 7(17), 6918–6926. 10.1002/ece3.3192

13. Ellison, A. M. (2010). Partitioning diversity. Ecology, 91(7), 1962–1963.

14. Faith, D. P. (1992). Conservation evaluation and phylogenetic diversity. Biological Conservation, 61, 1–10. 10.1016/0003-2697(75)90168-2

15. Fenoglio, M. S., Calviño, A., González, E., Salvo, A., & Videla, M. (2021). Urbanisation drivers and underlying mechanisms of terrestrial insect diversity loss in cities. Ecological Entomology, 46(4), 757–771. 10.1111/een.13041

16. Ghisbain, G., Thiery, W., Massonnet, F., Erazo, D., Rasmont, P., Michez, D., & Dellicour, S. (2024). Projected decline in European bumblebee populations in the twenty-first century. Nature, 628, 337–341. 10.1038/s41586-023-06471-0

17. Goslee, S., & Urban, D. (2007). The ecodist package for dissimilarity-based analysis of ecological data. Journal of Statistical Software, 22(7), 1–19.

18. Gotelli, N. J., & Colwell, R. K. (2001). Quantifying biodiversity: Procedures and pitfalls in the measurement and comparison of species richness. Ecology Letters, 4(4), 379–391. 10.1046/j.1461-0248.2001.00230.x

19. Gower, A. J. C. (1971). A general coefficient of similarity and some of its properties. Biometrics, 27(4), 857–871.

20. Hallmann, C. A., Sorg, M., Jongejans, E., Siepel, H., Hofland, N., Schwan, H., Stenmans, W., Müller, A., Sumser, H., Hörren, T., Goulson, D., & De Kroon, H. (2017). More than 75 percent decline over 27 years in total flying insect biomass in protected areas. PLoS ONE, 12(10), e0185809. 10.1371/journal.pone.0185809

21. Hallmann, C. A., Ssymank, A., Sorg, M., de Kroon, H., & Jongejans, E. (2021). Insect biomass decline scaled to species diversity: General patterns derived from a hoverfly community. Proceedings of the National Academy of Sciences of the United States of America, 118(2), 1–8. 10.1073/PNAS.2002554117

22. Hausmann, A., Segerer, A. H., Greifenstein, T., Knubben, J., Morinière, J., Bozicevic, V., Doczkal, D., Günter, A., Ulrich, W., & Habel, J. C. (2020). Toward a standardized quantitative and qualitative insect monitoring scheme. Ecology and Evolution, 10(9), 4009–4020. 10.1002/ece3.6166

23. Herrmann, J., Buchholz, S., & Theodorou, P. (2023). The degree of urbanisation reduces wild bee and butterfly diversity and alters the patterns of flower-visitation in urban dry grasslands. Scientific Reports, 13(1), 1–15. 10.1038/s41598-023-29275-8

24. Hill, M. O. (1973). Diversity and evenness: A unifying notation and its consequences. Ecology, 54(2), 427–432. 10.2307/1934352

25. Hothorn, T., Bretz, F., & Westfall, P. (2008). Simultaneous inference in general parametric models. Biometrical Journal, 50(3), 346–363. 10.1002/bimj.200810425

26. Knop, E. (2016). Biotic homogenization of three insect groups due to urbanization. Global Change Biology, 22(1), 228–236. 10.1111/gcb.13091

27. Macgregor, C. J., Williams, J. H., Bell, J. R., & Thomas, C. D. (2021). Moth biomass has fluctuated over 50 years in Britain but lacks a clear trend. Nature Ecology & Evolution, 3, 1645–1649. 10.1038/s41559-021-01449-5

28. Maechler, M., Rousseeuw, P., Struyf, A., Hubert, M., & Hornik, K. (2022). cluster: Cluster Analysis Basics and Extensions.

29. Moretti, M., Dias, A. T. C., de Bello, F., Altermatt, F., Chown, S. L., Azcárate, F. M., Bell, J. R., Fournier, B., Hedde, M., Hortal, J., Ibanez, S., Öckinger, E., Sousa, J. P., Ellers, J., & Berg, M. P. (2017). Handbook of protocols for standardized measurement of terrestrial invertebrate functional traits. Functional Ecology, 31(3), 558–567. 10.1111/1365-2435.12776

30. Müller, J., Hothorn, T., Yuan, Y., Seibold, S., Mitesser, O., Rothacher, J., Freund, J., Wild, C., Wolz, M., & Menzel, A. (2024). Weather explains the decline and rise of insect biomass over 34 years. Nature, 628, 349–354. 10.1038/s41586-023-06402-z

31. Neff, F., Korner-Nievergelt, F., Rey, E., Albrecht, M., Bollmann, K., Cahenzli, F., Chittaro, Y., Gossner, M. M., Martínez-Núñez, C., Meier, E. S., Monnerat, C., Moretti, M., Roth, T., Herzog, F., & Knop, E. (2022). Different roles of concurring climate and regional land-use changes in past 40 years’ insect trends. Nature Communications, 13(1), 11–13. 10.1038/s41467-022-35223-3

32. Nicholson, C. C., Knapp, J., Kiljanek, T., Albrecht, M., Chauzat, M. P., Costa, C., De la Rúa, P., Klein, A. M., Mänd, M., Potts, S. G., Schweiger, O., Bottero, I., Cini, E., de Miranda, J. R., Di Prisco, G., Dominik, C., Hodge, S., Kaunath, V., Knauer, A., … Rundlöf, M. (2023). Pesticide use negatively affects bumble bees across European landscapes. Nature, 628, 355–358. 10.1038/s41586-023-06773-3

33. Outhwaite, C. L., McCann, P., & Newbold, T. (2022). Agriculture and climate change are reshaping insect biodiversity worldwide. Nature, 605(7908), 97–102. 10.1038/s41586-022-04644-x

34. Paradis, E., & Schliep, K. (2019). Ape 5.0: An environment for modern phylogenetics and evolutionary analyses in R. Bioinformatics, 35(3), 526–528. 10.1093/bioinformatics/bty633

35. Pilotto, F., Kühn, I., Adrian, R., Alber, R., Alignier, A., Andrews, C., Bäck, J., Barbaro, L., Beaumont, D., Beenaerts, N., Benham, S., Boukal, D. S., Bretagnolle, V., Camatti, E., Canullo, R., Cardoso, P. G., Ens, B. J., Everaert, G., Evtimova, V., … Haase, P. (2020). Meta-analysis of multidecadal biodiversity trends in Europe. Nature Communications, 11, 3486. 10.1038/s41467-020-17171-y

36. Pont, D., Meulenbroek, P., Bammer, V., Dejean, T., Erős, T., Jean, P., Lenhardt, M., Nagel, C., Pekarik, L., Schabuss, M., Stoeckle, B. C., Stoica, E., Zornig, H., Weigand, A., & Valentini, A. (2023). Quantitative monitoring of diverse fish communities on a large scale combining eDNA metabarcoding and qPCR. Molecular Ecology Resources, 23(2), 396–409. 10.1111/1755-0998.13715

37. Powney, G. D., Carvell, C., Edwards, M., Morris, R. K. A., Roy, H. E., Woodcock, B. A., & Isaac, N. J. B. (2019). Widespread losses of pollinating insects in Britain. Nature Communications, 10, 1018. 10.1038/s41467-019-08974-9

38. R Core Team. (2023). R: A Language and Environment for Statistical Computing. R Foundation for Statistical Computing. https://www.r-project.org/

39. Rainford, J. L., Hofreiter, M., Nicholson, D. B., & Mayhew, P. J. (2014). Phylogenetic distribution of extant richness suggests metamorphosis is a key innovation driving diversification in insects. PLoS ONE, 9(10), e109085. 10.1371/journal.pone.0109085

41. Raven, P. H., & Wagner, D. L. (2021). Agricultural intensification and climate change are rapidly decreasing insect biodiversity. PNAS, 118(2), e2002548117. 10.1073/PNAS.2002548117

42. Redlich, S., Zhang, J., Benjamin, C., Dhillon, M. S., Englmeier, J., Ewald, J., Fricke, U., Ganuza, C., Haensel, M., Hovestadt, T., Kollmann, J., Koellner, T., Kübert-Flock, C., Kunstmann, H., Menzel, A., Moning, C., Peters, W., Riebl, R., Rummler, T., … Steffan-Dewenter, I. (2022). Disentangling effects of climate and land use on biodiversity and ecosystem services – a multi- scale experimental design. Methods in Ecology and Evolu় on, 13, 514–527. 10.1111/2041-210X.13759

43. Roswell, M., Dushoff, J., & Winfree, R. (2021). A conceptual guide to measuring species diversity. Oikos, 130(3), 321–338. 10.1111/oik.07202

44. Saunders, M. E., Janes, J. K., & O’Hanlon, J. C. (2020). Moving on from the Insect Apocalypse Narrative: Engaging with Evidence-Based Insect Conservation. BioScience, 70(1), 80–89. 10.1093/biosci/biz143

45. Savage, A. M., Hackett, B., Guénard, B., Youngsteadt, E. K., & Dunn, R. R. (2015). Fine-scale heterogeneity across Manhattan’s urban habitat mosaic is associated with variation in ant composition and richness. Insect Conservation and Diversity, 8(3), 216–228. 10.1111/icad.12098

46. Seibold, S., Brandl, R., Buse, J., Hothorn, T., Schmidl, J., Thorn, S., & Müller, J. (2015). Association of extinction risk of saproxylic beetles with ecological degradation of forests in Europe. Conservation Biology, 29(2), 382–390. 10.1111/cobi.12427

47. Seibold, S., Gossner, M. M., Simons, N. K., Blüthgen, N., Müller, J., Ambarlı, D., Ammer, C., Bauhus, J., Fischer, M., Habel, J. C., Linsenmair, K. E., Nauss, T., Penone, C., Prati, D., Schall, P., Schulze, E. D., Vogt, J., Wöllauer, S., & Weisser, W. W. (2019). Arthropod decline in grasslands and forests is associated with landscape-level drivers. Nature, 574(7780), 671–674. 10.1038/s41586-019-1684-3

48. Staab, M., Gossner, M. M., Simons, N. K., Achury, R., Ambarlı, D., Bae, S., Schall, P., Weisser, W. W., & Blüthgen, N. (2023). Insect decline in forests depends on species’ traits and may be mitigated by management. Communications Biology, 6(1), 1–13. 10.1038/s42003-023-04690-9

49. Svenningsen, C. S., Bowler, D. E., Hecker, S., Bladt, J., Grescho, V., van Dam, N. M., Dauber, J., Eichenberg, D., Ejrnæs, R., Fløjgaard, C., Frenzel, M., Frøslev, T. G., Hansen, A. J., Heilmann-Clausen, J., Huang, Y., Larsen, J. C., Menger, J., Nayan, N. L. B. M., Pedersen, L. B., … Bonn, A. (2022). Flying insect biomass is negatively associated with urban cover in surrounding landscapes. Diversity and Distributions, 28(6), 1242–1254. 10.1111/ddi.13532

50. Uhler, J., Haase, P., Hoffmann, L., Hothorn, T., Schmidl, J., Stoll, S., Welti, E. A. R., Buse, J., & Müller, J. (2022). A comparison of different Malaise trap types. Insect Conservation and Diversity, 15(6), 666–672. 10.1111/icad.12604

51. Uhler, J., Redlich, S., Zhang, J., Hothorn, T., Tobisch, C., Ewald, J., Thorn, S., Seibold, S., Mitesser, O., Morinière, J., Bozicevic, V., Benjamin, C. S., Englmeier, J., Fricke, U., Ganuza, C., Haensel, M., Riebl, R., Rojas-Botero, S., Rummler, T., … Müller, J. (2021). Relationship of insect biomass and richness with land use along a climate gradient. Nature Communications, 12, 5946. 10.1038/s41467-021-26181-3

52. van Klink, R., Bowler, D. E., Gongalky, K. B., Swengel, A. B., Gentiel, A., & Chase, J. M. (2020). Meta-analysis reveals declines in terrestrial but increases in freshwater insect abundances. Science, 368, 417–420. 10.1126/science.abd8947

53. van Klink, R., Bowler, D. E., Gongalsky, K. B., Shen, M., Swengel, S. R., & Chase, J. M. (2024). Disproportionate declines of formerly abundant species underlie insect loss. Nature, 628, 359–364. 10.1038/s41586-023-06861-4

54. Wagner, D. L. (2020). Insect declines in the anthropocene. Annual Review of Entomology, 65, 457–480. 10.1146/annurev-ento-011019-025151

55. Webb, C. O., Ackerly, D. D., & Kembel, S. W. (2008). Phylocom: Software for the analysis of phylogenetic community structure and trait evolution. Bioinformatics, 24(18), 2098–2100. 10.1093/bioinformatics/btn358

56. Winter, M., Devictor, V., & Schweiger, O. (2013). Phylogenetic diversity and nature conservation: Where are we? Trends in Ecology and Evolution, 28(4), 199–204. 10.1016/j.tree.2012.10.015

57. Wright, E. S. (2015). DECIPHER: Harnessing local sequence context to improve protein multiple sequence alignment. BMC Bioinformatics, 16(1), 1–14. 10.1186/s12859-015-0749-z

