## Supplement 1 for "A short cut to sample coverage standardization in meta-barcoding data provides new insights into land use effects on insect diversity"

**Fig. S1.1. Overview of the statistical methods.** The numbers in the blue boxes refer to the R scripts in the Zenodo repository https://doi.org/10.5281/zenodo.14747928.

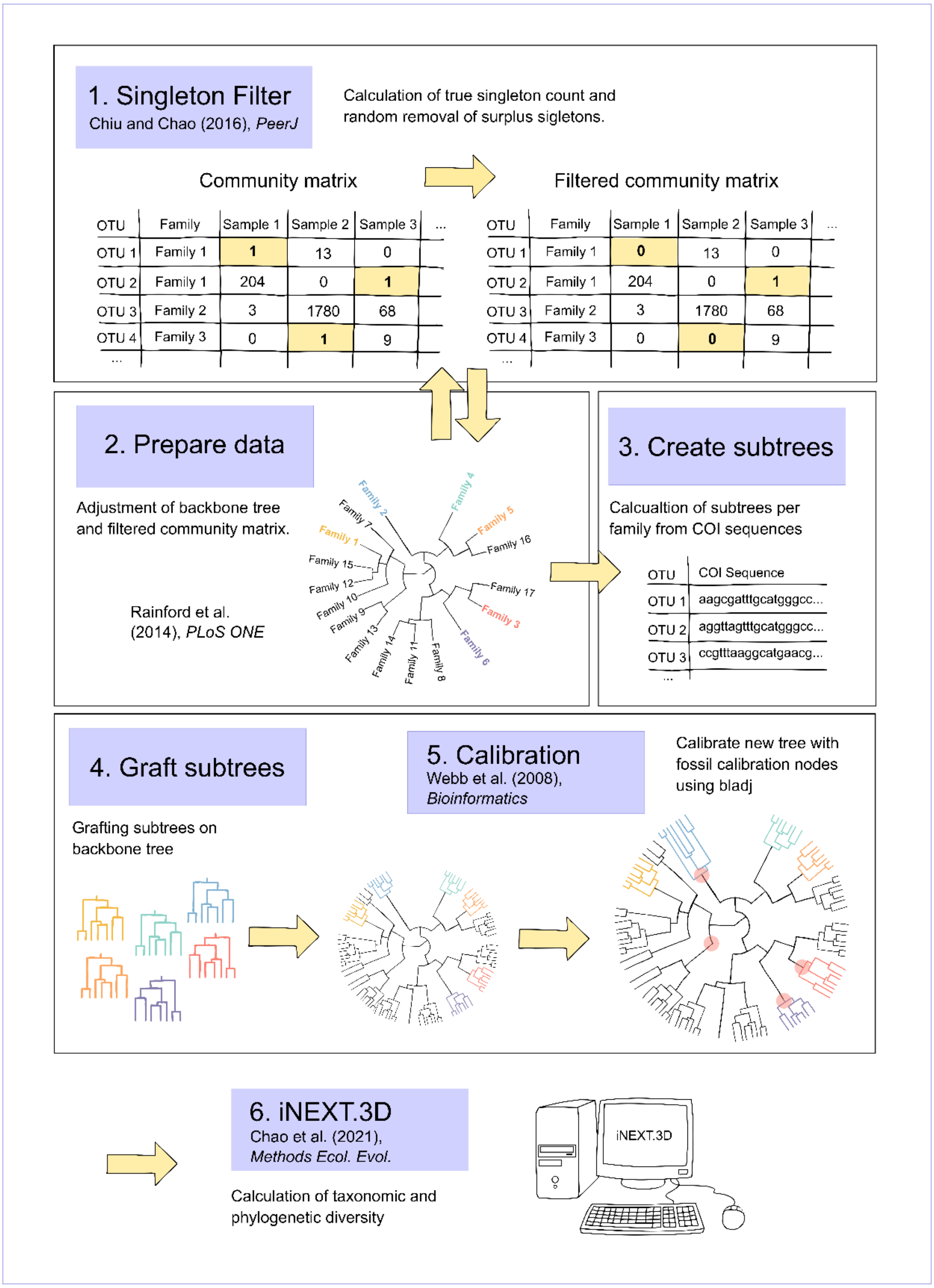

**Fig. S1.2. Histogram of the sample coverage of the analysed insect samples.**

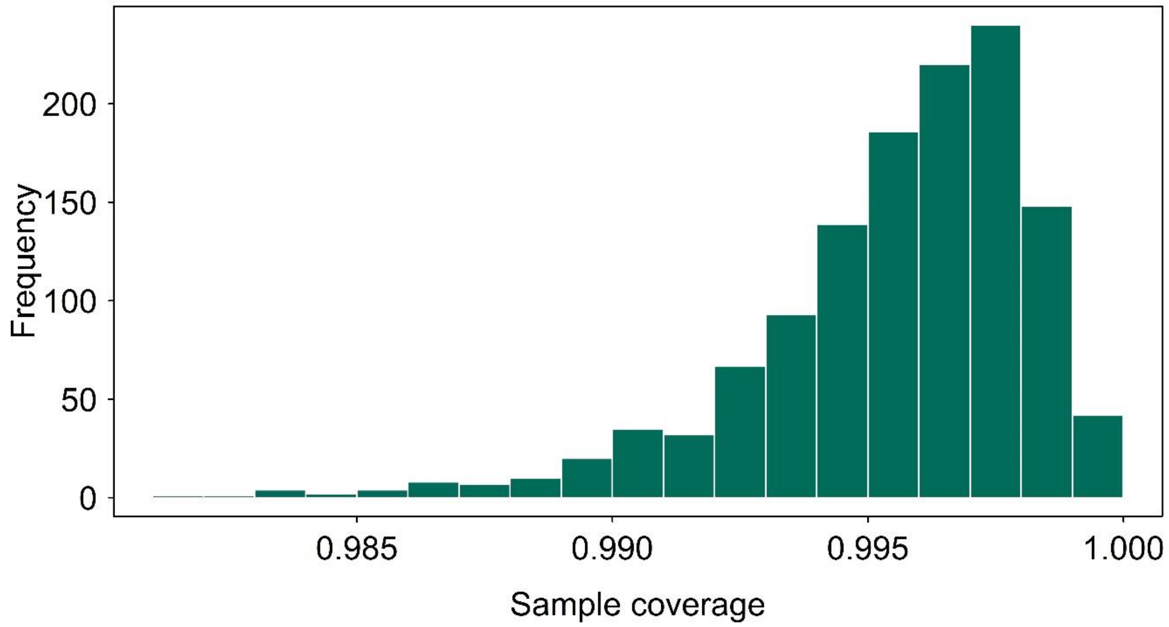


**Fig. S1.3. Multiplicative effects of season on sample coverage of insects.** Partial effects from generalized additive mixed models were controlled for elevation, the geographic location of the Malaise traps, and local and regional land-use types (for details, see Table S1).

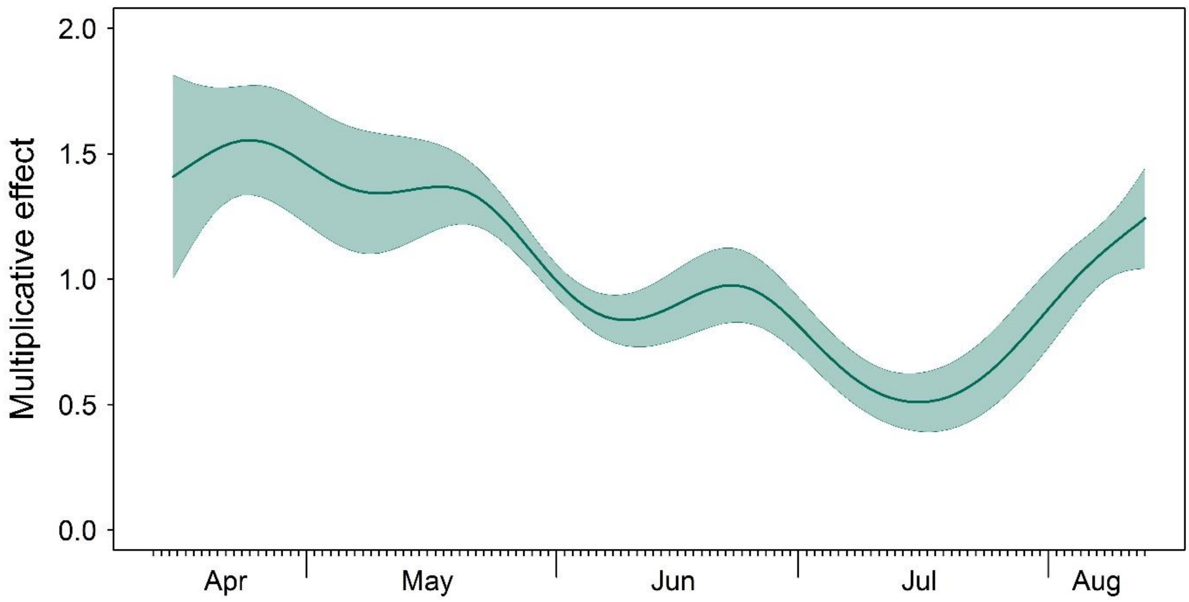

**Table S1.1. Results of generalized additive mixed models including local and regional land-use categories.** Taxonomic and phylogenetic diversity were calculated with focus on rare (q=0), common (q=1) and dominant (q=2) species and for a standardized sample coverage of 99.6%. The model included local land-use types (arable land, grassland and settlement, with forest as base) and regional land-use types (agricultural and urban with semi-natural as base) climate (long-term precipitation and long-term temperature) and weather (temperature and humidity) conditions, mean day of the year and geographic location (coordinates). Significant results are depicted in bold.

| | Sample Coverage | | Taxonomic diversity | | | | | | Phylogenetic diversity | | | | | |
|---|---|---|---|---|---|---|---|---|---|---|---|---|---|---|
| | | | q=0 | | q=1 | | q=2 | | q=0 | | q=1 | | q=2 | |
| | Estimate | p.value | Estimate | p.value | Estimate | p.value | Estimate | p.value | Estimate | p.value | Estimate | p.value | Estimate | p.value |
| Intercept | 3.159 | **<0.001** | 2.168 | **<0.001** | -0.080 | 0.817 | -1.035 | **0.006** | 1.164 | **<0.001** | -2.172 | **<0.001** | -2.516 | **<0.001** |
| Arable land | 0.106 | **0.007** | -0.235 | **<0.001** | -0.349 | **<0.001** | -0.317 | **<0.001** | -0.204 | **<0.001** | -0.202 | **<0.001** | -0.093 | **<0.001** |
| Grassland | -0.002 | 0.959 | -0.094 | **0.001** | -0.183 | **<0.001** | -0.181 | **<0.001** | -0.081 | **<0.001** | -0.095 | **<0.001** | -0.033 | **0.011** |
| Settlement | -0.063 | 0.133 | -0.050 | 0.142 | -0.128 | **0.001** | -0.117 | **0.007** | -0.058 | **0.026** | -0.103 | **<0.001** | -0.037 | **0.013** |
| Agricultural | 0.033 | 0.338 | -0.099 | **<0.001** | -0.154 | **<0.001** | -0.140 | **<0.001** | -0.092 | **<0.001** | -0.108 | **<0.001** | -0.057 | **<0.001** |
| Urban | 0.121 | **<0.001** | -0.079 | **0.004** | -0.095 | **0.003** | -0.075 | **0.033** | -0.064 | **0.003** | -0.035 | **0.040** | -0.025 | **0.043** |
| Long-term precipitation | <0.001 | 0.484 | <0.001 | 0.148 | <0.001 | **0.039** | <0.001 | **0.038** | <0.001 | 0.123 | <0.001 | **<0.001** | 0.000 | **<0.001** |
| Long-term temperature | -0.047 | 0.063 | 0.048 | **0.015** | 0.051 | **0.032** | 0.039 | 0.130 | 0.013 | 0.393 | 0.018 | 0.153 | 0.003 | 0.776 |
| Temperature | -0.003 | 0.820 | 0.057 | **<0.001** | 0.069 | **<0.001** | 0.073 | **<0.001** | 0.049 | **<0.001** | 0.046 | **<0.001** | 0.032 | **<0.001** |
| Humidity | 0.001 | 0.834 | 0.004 | 0.077 | 0.006 | **0.034** | 0.006 | **0.049** | 0.006 | **0.003** | 0.008 | **<0.001** | 0.003 | **0.004** |
| Mean day of the year | 8.037 | **<0.001** | 8.551 | **<0.001** | 8.675 | **<0.001** | 8.595 | **<0.001** | 8.252 | **<0.001** | 8.639 | **<0.001** | 8.319 | **<0.001** |
| Coordinates | 0.005 | 0.881 | 3.832 | 0.084 | 5.385 | **0.028** | 2.034 | 0.206 | 3.147 | 0.051 | 1.348 | **0.187** | 2.094 | 0.080 |

**Table S1.2. Results of generalized additive mixed models including local and regional land-use categories.** Taxonomic and phylogenetic diversity were calculated with focus on rare (q=0), common (q=1) and dominant (q=2) species for a standardized sample coverage of 98%. The model included local land-use types (arable land, grassland and settlement, with forest as base) and regional land-use types (agricultural and urban with semi-natural as base) climate (long-term precipitation and long-term temperature) and weather (temperature and humidity) conditions, mean day of the year and geographic location (coordinates). Significant results are depicted in bold.

| | Sample Coverage | | Taxonomic diversity | | | | | | Phylogenetic diversity | | | | | |
|---|---|---|---|---|---|---|---|---|---|---|---|---|---|---|
| | | | q=0 | | q=1 | | q=2 | | q=0 | | q=1 | | q=2 | |
| | Estimate | p.value | Estimate | p.value | Estimate | p.value | Estimate | p.value | Estimate | p.value | Estimate | p.value | Estimate | p.value |
| Intercept | 3.159 | **<0.001** | 1.875 | **<0.001** | -0.096 | 0.783 | -1.049 | **0.005** | 0.671 | **0.008** | -2.181 | **<0.001** | -2.517 | **<0.001** |
| Arable land | 0.106 | **0.007** | -0.261 | **<0.001** | -0.349 | **<0.001** | -0.317 | **<0.001** | -0.240 | **<0.001** | -0.202 | **<0.001** | -0.093 | **<0.001** |
| Grassland | -0.002 | 0.959 | -0.103 | **0.001** | -0.182 | **<0.001** | -0.180 | **<0.001** | -0.090 | **<0.001** | -0.095 | **<0.001** | -0.033 | **0.011** |
| Settlement | -0.063 | 0.133 | -0.058 | 0.097 | -0.127 | **0.001** | -0.117 | **0.007** | -0.072 | **0.013** | -0.103 | **<0.001** | -0.037 | 0.013 |
| Agricultural | 0.033 | 0.338 | -0.111 | **<0.001** | -0.155 | **<0.001** | -0.140 | **<0.001** | -0.113 | **<0.001** | -0.108 | **<0.001** | -0.057 | **<0.001** |
| Urban | 0.121 | **<0.001** | -0.084 | **0.003** | -0.095 | **0.003** | -0.076 | **0.030** | -0.062 | **0.010** | -0.035 | **0.039** | -0.025 | 0.043 |
| Long-term precipitation | <0.001 | 0.484 | <0.001 | 0.124 | <0.001 | **0.037** | <0.001 | **0.038** | 0.000 | 0.174 | 0.000 | **<0.001** | 0.000 | **<0.001** |
| Long-term temperature | -0.047 | 0.063 | 0.056 | **0.008** | 0.051 | **0.032** | 0.039 | 0.122 | 0.027 | 0.131 | 0.018 | 0.151 | 0.003 | 0.775 |
| Temperature | -0.003 | 0.820 | 0.055 | **<0.001** | 0.068 | **<0.001** | 0.073 | **<0.001** | 0.046 | **<0.001** | 0.046 | **<0.001** | 0.033 | **<0.001** |
| Humidity | 0.001 | 0.834 | 0.005 | 0.064 | 0.006 | **0.034** | 0.006 | **0.048** | 0.007 | **0.002** | 0.008 | **<0.001** | 0.003 | **0.004** |
| Mean day of the year | 8.037 | **<0.001** | 8.582 | **<0.001** | 8.675 | **<0.001** | 8.597 | **<0.001** | 8.438 | **<0.001** | 8.640 | **<0.001** | 8.319 | **<0.001** |
| Coordinates | 0.005 | 0.881 | 7.307 | **0.017** | 5.568 | **0.026** | 2.113 | 0.198 | 14.521 | **0.002** | 1.357 | 0.186 | 2.095 | 0.080 |
